# Sleep modulates neural timescales and spatiotemporal integration in the human cortex

**DOI:** 10.1101/2024.09.26.614972

**Authors:** Riccardo Cusinato, Andrea Seiler, Kaspar Schindler, Athina Tzovara

## Abstract

Spontaneous neural dynamics manifest across multiple timescales, which are intrinsic to brain areas and exhibit hierarchical organization across the cortex. In wake, a hierarchy of timescales is thought to naturally emerge from microstructural properties, gene expression, and recurrent connections. A fundamental question is timescales’ organization and changes in sleep, where physiological needs are different. Here, we describe two coexisting but distinct measures of neural timescales, obtained from broadband activity and gamma power, which display complementary properties. We leveraged intracranial electroencephalography (iEEG) data to characterize timescale changes from wake to sleep across the cortical hierarchy. We show that both broadband and gamma timescales are globally longer in sleep than in wake. While broadband timescales increase along the sensorimotor-association axis, gamma ones decrease. During sleep, slow waves can explain the increase of broadband and gamma timescales, but only broadband ones show a positive association with slow-wave density across the cortex. Finally, we characterize spatial correlations and their relationship with timescales as a proxy for spatiotemporal integration, finding high integration at long distances in wake for broadband and at short distances in sleep for gamma timescales. Our results suggest that mesoscopic neural populations possess different timescales that are shaped by anatomy and are modulated by the sleep/wake cycle.

**Significance statement:** Understanding the organization of intrinsic neural dynamics is crucial for investigating brain functions in health and disease. A key question is: how do neural dynamics change in the sleeping brain? Here we focus on neural timescales, which measure temporal autocorrelation and are organized hierarchically across the cortex, and spatial correlations. We show that two types of timescales exist in neural populations recorded with intracranial electroencephalography in humans, corresponding to broadband (0.5-80 Hz) and gamma (40-80 Hz) frequency ranges. Both timescales increase in sleep, where slow waves have an important role, but follow opposite hierarchies: broadband timescales increase from sensory to associative areas, while gamma timescales show the reverse pattern. Finally, timescales covary with spatial correlations, suggesting higher spatiotemporal integration over long distances in wake compared to sleep.

## Introduction

The study of intrinsic neural dynamics is fundamental for understanding brain function. Neural activity fluctuates at a characteristic timing, the neural timescale, which follows a hierarchical organization across measurement modalities and species (Cusinato et al., 2023; A. M. Manea et al., 2022; Murray et al., 2014; Raut et al., 2020). Neural timescales are short in sensory areas and progressively increase while advancing through cortical hierarchies (Chaudhuri et al., 2015; Gao et al., 2020; Murray et al., 2014). This regional specificity can be explained by the underlying microstructure, including myelination content (Burt et al., 2018), expression of excitation/inhibition-related genes (Gao et al., 2020; Shafiei et al., 2023), or connectivity patterns across- and within-areas (Stern et al., 2023). Strongly interconnected brain areas show longer timescales than weakly connected ones (Chaudhuri et al., 2015; Fallon et al., 2020), consistent with the hypothesized role of timescales in integrating information (Golesorkhi et al., 2021).

The prevalent view on neural timescales is that they are intrinsic properties of brain circuits (Murray et al., 2014; Rossi-Pool et al., 2021). Although the notion of “intrinsic” implies that timescales are hard-wired, single-neuron timescales display considerable variability already within a single area (Cavanagh et al., 2020) and spiking activity exhibits multiple timescales (Zeraati et al., 2023). Additionally, neural oscillations in cortical areas manifest at different characteristic frequencies, which are hierarchically organized but not unique, raising the possibility of coexisting timescales within single areas (Vinck et al., 2023).

Neural timescales also change with brain states (Cavanagh et al., 2020), particularly when transitioning from wake to sleep (Müller & Meisel, 2023; Zilio et al., 2021). In sleep, neural dynamics are drastically altered (Luppi et al., 2024), as global oscillatory signatures are visible in the EEG (Adamantidis et al., 2019). In humans, rapid eye movement (REM) and non-REM 3 (NREM3) sleep are primarily characterized by theta oscillations and slow waves, respectively (Cantero et al., 2003; Massimini et al., 2004). However, sleep dynamics are not global and exhibit local changes across cortical areas in a range of frequencies (Andrillon et al., 2011; Nir et al., 2011). It follows naturally that the well-described sensorimotor-association hierarchy of fast-to-slow neural timescales in the awake brain may be reorganized during sleep, but existing studies have mixed conclusions. In humans, timescales have been found to globally increase during sleep when measured via scalp EEG (Zilio et al., 2021), functional magnetic resonance imaging (fMRI) (Klar et al., 2023), and spiking activity (Hagemann et al., 2022), but to decrease when measured from gamma power in iEEG (Müller & Meisel, 2023). One explanation for this discrepancy might be related to the heterogeneity of timescales in the awake brain, which change non-uniformly in fine-grained circuits when transitioning to sleep.

Sleep is also a state of altered connectivity between cortical areas, with NREM sleep traditionally seen as a “disconnected” stage in which neural communication is hampered, and REM as more similar to wake (Massimini et al., 2005, 2010). In wake, areas with higher timescales show stronger functional and structural connectivity (Fallon et al., 2020; Lurie et al., 2024; Shinn et al., 2023). This highlights the importance of quantifying not only temporal but also spatial correlations to understand large-scale cortical organization and spatiotemporal integration.

Here, we sought to characterize neural timescales, spatial correlations, and their relationship in wake and sleep. We hypothesized a link between timescales and spatial correlations in human iEEG, with a regional reconfiguration of both measures in sleep; and that iEEG neural activity would exhibit an overlapping variety of timescales related to different frequency bands. We identified two hierarchies of neural timescales, derived from the broadband iEEG and the gamma power signals. They both increased in sleep compared to wake and were similarly affected by slow-wave events but showed an opposite relationship with the sensorimotor-association hierarchy. Finally, we also showed that timescales and spatial correlations are related, but at different spatial scales and in a stage-dependent manner.

## Results

To understand the organization of neural timescales and spatial correlations (SC) across the cortex in wake and sleep, we analyzed resting-state intracranial EEG (iEEG) of a large cohort of patients from the MNI open iEEG dataset (106 patients, 48 females, 13–62 years old) (Figure 1). We utilized 1-minute recordings during wake, NREM3 and REM sleep, from 1772 contacts (Figure 1A, in wake, reduced to 1468 in NREM3 and 1012 in REM sleep; Figure 1B, exemplar signals in a temporal and a frontal channel).

**Figure 1.**
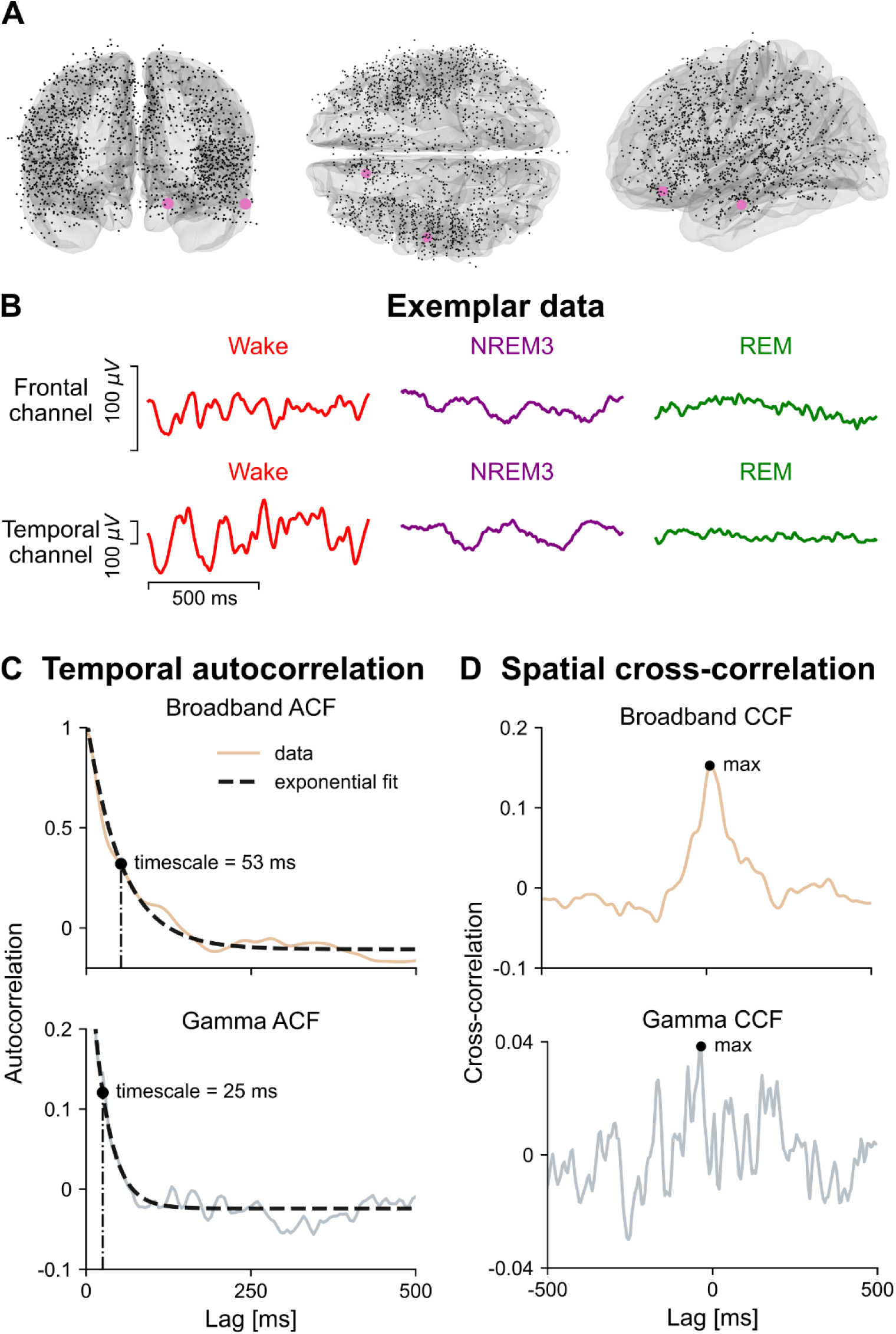
Overview of dataset and methods. A) Locations of the 1772 contacts in the MNI Open iEEG dataset, from frontal, superior and lateral views in MNI space. The contacts shown are those with wake data, while recorded contacts in sleep are less (1468 for NREM3 and 1012 for REM). Highlighted are two exemplar contacts in the frontal and temporal lobes. B) Representative 1-second iEEG traces from the two contacts, in Wake, NREM3 and REM. Sleep stages exhibit characteristic features, i.e., slow high-amplitude fluctuations in NREM3 and desynchronized low-amplitude fluctuations in REM. Note that the y-axis scale is different for the two contacts. Time series are processed as “broadband” (0.5-80 Hz voltage, i.e. iEEG) and “gamma” (40-80 Hz power fluctuations). C) Both signals from each contact and each stage are used to compute the autocorrelation function (ACF) and the related exponential fit. The model is of the form *ACF*(*k*) = *a*(*e*^−*k*/*τ*^ + *b*), with *τ* being the timescale. Here just one exemplar ACF is shown. D) The cross-correlation function (CCF) is also computed for the broadband and gamma signals, between pairs of channels. From it, the maximum cross-correlation across lags is used.

We quantified broadband and gamma timescales at the mesoscale level via an exponential fit to the autocorrelation function (ACF) (Figure 1C). We also computed a measure of SC, specifically the maximum cross-correlation function (CCF) across lags between pairs of channels (Figure 1D).

### Broadband timescales increase in sleep and along the sensorimotor-association hierarchy

The average broadband ACF exhibited higher autocorrelations in NREM3 sleep and a moderate increase in REM sleep (Figure 2A). To parcellate iEEG channels into cortical areas, we used the 180 left-hemisphere areas of the HCP-MMP parcellation. All cortical areas showed a marked increase in broadband timescales from wake to NREM3 (mean increase 105 ms, [102, 108] 95% CI with bootstrap) and 163/180 areas showed a moderate increase in REM sleep (mean increase 16 ms, [14, 18] 95% CI with bootstrap). Only one area showed an increase from NREM3 to REM sleep, the presubiculum in the medial temporal lobe (MTL) (Figure 2B). In NREM3 the distribution of broadband timescales was quite uniform (Figure 2C), with lower values in medial temporal lobe, auditory cortex, occipital and medial regions. In REM, there was a marked increase of timescales in the temporal lobe, especially in MTL, auditory cortex and temporal pole.

**Figure 2.**
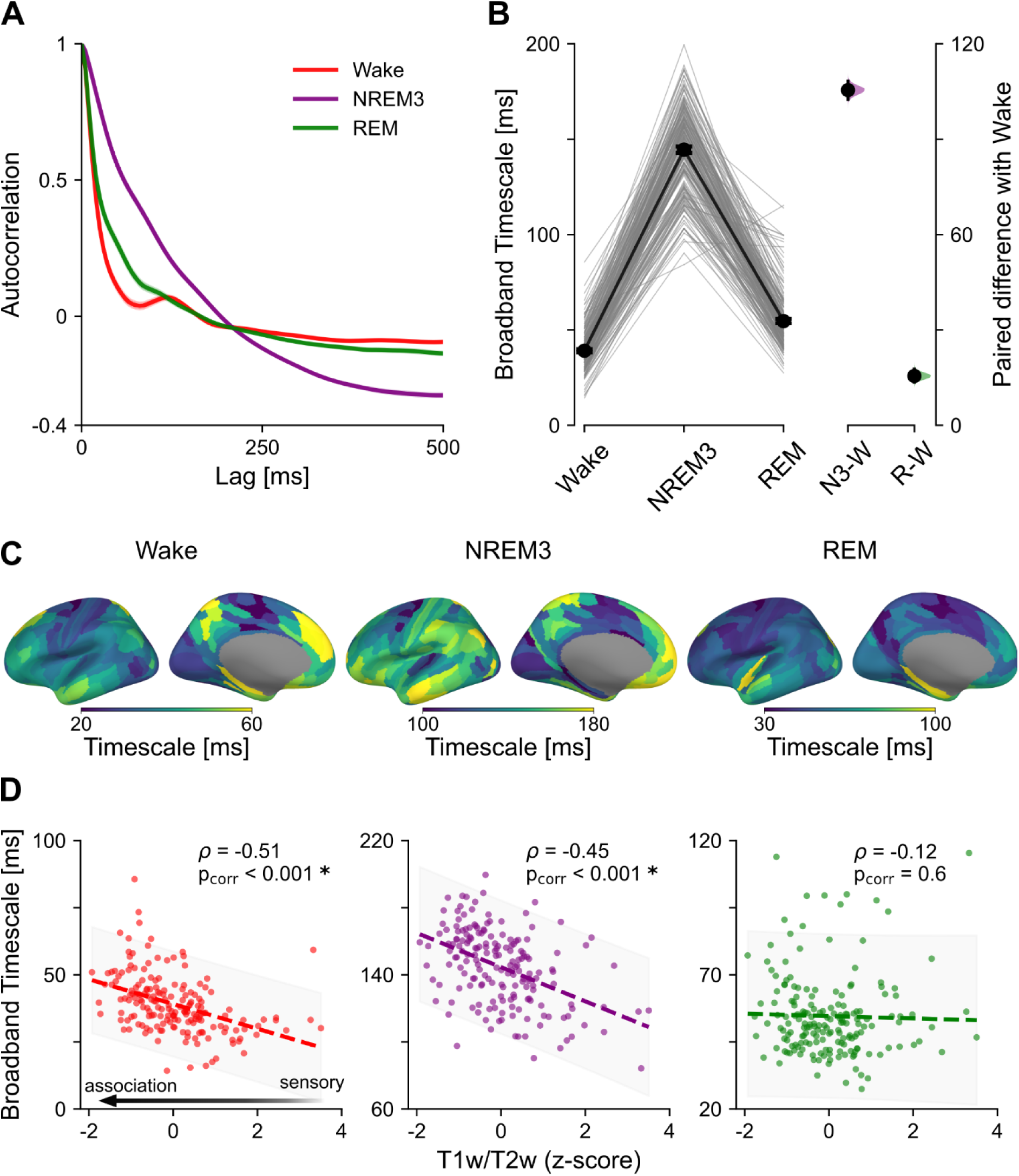
Broadband timescales across stages and relation to the anatomical hierarchy. A) Average ACF for wake, NREM3 and REM, averaged first across channels in a patient and then across patients. Shaded areas represent s.e.m. across patients. Note how autocorrelations in NREM3 remain higher at longer lags than in the other stages. B) Broadband timescales were parcellated in the 180 areas of the HCP-MMP parcellation and projected to the left hemisphere. Left: Slope plot with the change of broadband timescales in all cortical areas across stages; thin lines represent each area and black dots average values. Right: Estimation plot with the effect size of the difference between sleep stages and wake. The central dot represents the mean value and the vertical bar the 95% confidence interval; distributions are obtained via 9999 bootstraps. C) Broadband timescales projected onto the “fsaverage” inflated surface template, with brighter colors indicating higher values. The values in sleep change drastically from wake. Note the different scales of the colormaps. D) Correlation between broadband timescales and z-scored T1w/T2w, for each sleep stage. The T1w/T2w is a measure of anatomical cortical hierarchy, and higher values indicate more sensory regions. The correlation is computed with Spearman’s correlation coefficient and p-values are corrected for spatial autocorrelation via permutations with the “vasa” method. Shaded areas represent the 95% prediction interval of the regression. Note the different scales of the y-axes. s.e.m. = standard error of the mean.

The distribution of broadband timescales across cortical areas revealed a precise relationship with the anatomical hierarchy. As previously reported in (Gao et al., 2020), broadband timescales in wake increased along the sensorimotor-association axis, with the longest timescales in MTL, anterior cingulate cortex and orbitofrontal cortex. We used the T1w/T2w as a marker of myelination to define sensory and association areas along a continuum (Burt et al., 2018). We first confirmed the sensorimotor-association increase in wake (ρ=-0.51, p_corr_<0.001) (Gao et al., 2020). During sleep, we found a similar sensorimotor-association increase of timescales in NREM3 (ρ=-0.45, p_corr_<0.001), but not in REM sleep (ρ=-0.12, p_corr_=0.60; all computed with Spearman’s coefficient and corrected for spatial autocorrelation with “vasa” method) (Figure 2D). The absence of hierarchy in REM could be attributed to the high value of broadband timescales in auditory and visual areas (Figure 3A).

**Figure 3.**
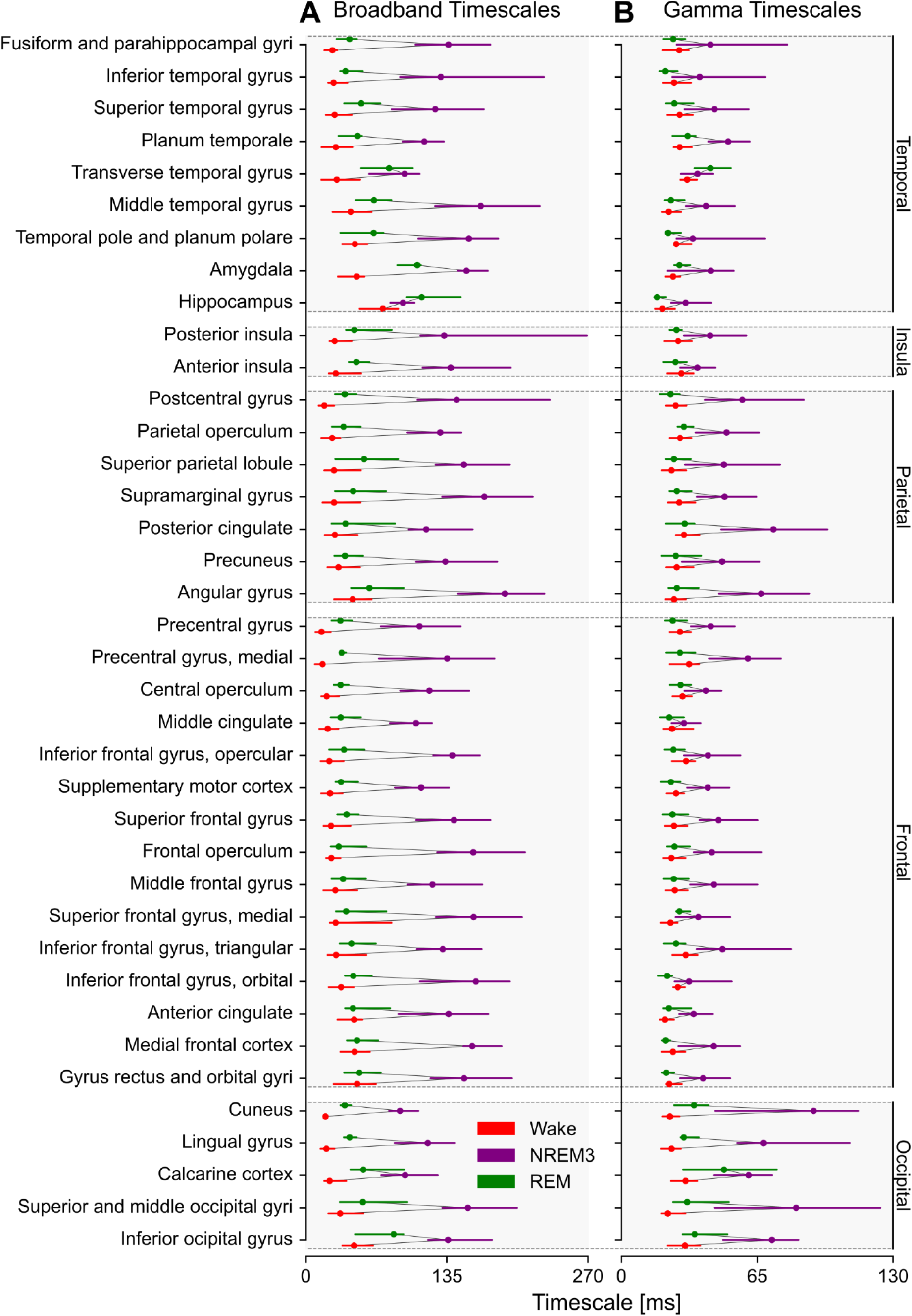
Overview of broadband and gamma timescales across the 38 areas of the MNI parcellation. Median timescale values together with the [25, 75] quantile range for broadband (A) and gamma power (B) signals. The areas are organized by lobe, which is written on the right. The areas’ order within lobe is given by increasing broadband timescale in wake and the order for gamma mirrors that of broadband. Note how all areas display increased timescales in sleep.

### Gamma timescales increase in sleep but decrease along the sensorimotor-association hierarchy

For gamma timescales, we found similar ACF profiles as for broadband ones (Figure 4A). However, autocorrelations in gamma power were generally weaker than broadband ones (compare y-axes of Figure 2A and Figure 4A). As for broadband, all cortical areas increased their gamma timescales in NREM3 (mean increase 31 ms, [29, 34] 95% CI with bootstrap), but just 92/180 had a moderate increase in REM (mean increase 2 ms, [1, 3] 95% CI with bootstrap) (Figure 4B). Interestingly, two areas increased their gamma timescale from NREM3 to REM: areas V6 and V7 in the occipital lobe. In addition to autocorrelation values, gamma timescales were lower than broadband ones (compare y-axes of Figure 2B and Figure 4B). We tested for a putative relationship between broadband and gamma timescales across areas but we didn’t find an obvious one for any stage (wake: ρ=-0.10, p_corr_=0.47; NREM3: ρ=-0.09, p_corr_=0.60; REM: ρ=0.01, p_corr_=0.99; all computed with Spearman’s coefficient and corrected for spatial autocorrelation with “vasa” method), suggesting they were not trivially related within individual areas (Figure 3).

**Figure 4.**
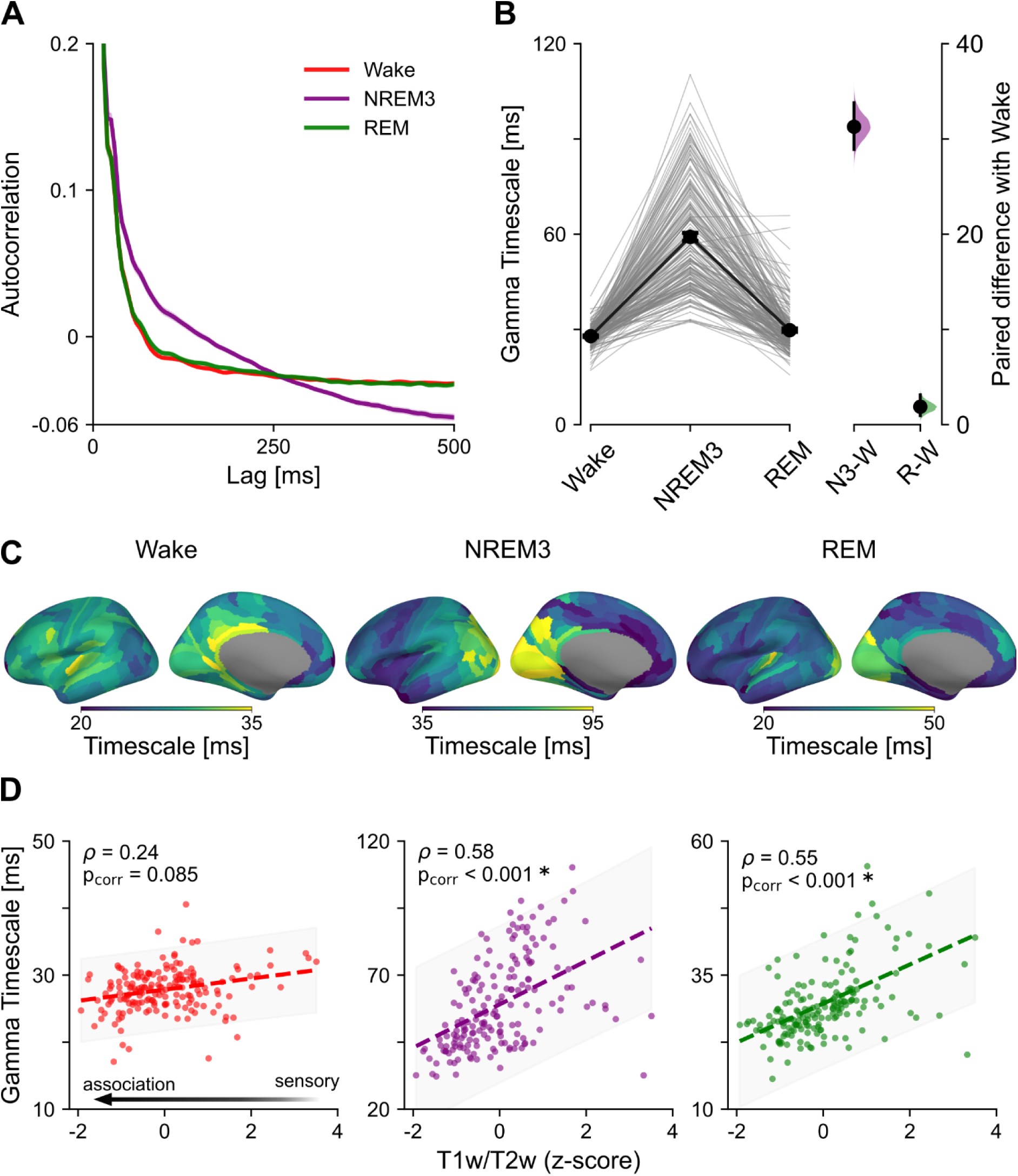
Gamma timescales across sleep stages and relation to the anatomical hierarchy. A) Average ACF for wake, NREM3 and REM, averaged first across channels in a patient and then across patients. Shaded areas represent s.e.m. across patients. Note how autocorrelations in NREM3 remain higher at longer lags than in the other stages. B) Gamma timescales were parcellated in the 180 areas of the HCP-MMP parcellation and projected to the left hemisphere. Left: Slope plot with the change of gamma timescales in all cortical areas across stages; thin lines represent each area and black dots average values. Right: Estimation plot with the effect size of the difference between sleep stages and wake. The central dot represents the mean and the vertical bar the 95% confidence interval; distributions are obtained via 9999 bootstraps. C) Gamma timescales projected onto the “fsaverage” inflated surface template, with brighter colors indicating higher values. Gamma timescales in sleep are generally higher in sensory regions. Note the different scales of the colormaps. D) Correlation between gamma timescales and z-scored T1w/T2w, for each sleep stage. The T1w/T2w is a measure of anatomical cortical hierarchy, and higher values indicate more sensory regions. The correlation is computed with Spearman’s correlation coefficients and p-values are corrected for spatial autocorrelation via permutations with the “vasa” method. Shaded areas represent the 95% prediction interval of the regression. Note the different scales of the y-axes. s.e.m. = standard error of the mean.

The distribution of gamma timescales across the cortex was markedly different from that of broadband ones (Figure 4C). In wake, higher gamma timescales were found in posterior cingulate and auditory areas; in NREM3 there was a prominent rostrocaudal gradient, with higher timescales in occipital regions; in REM, visual and auditory areas possessed the highest timescales (Figure 3B for a detailed overview). Contrary to broadband timescales, gamma ones were higher in sensory regions compared to associative ones, as reflected in a positive correlation with T1w/T2w (Figure 4D). The effect did not reach significance in wake following the strict correction for spatial autocorrelation, but was prominent in NREM3 and REM sleep (wake: ρ=0.24, p_corr_=0.09; NREM3: ρ=0.58, p_corr_<0.001; REM: ρ=0.55, p_corr_<0.001; all computed with Spearman’s coefficient and corrected for spatial autocorrelation with “vasa” method).

These results indicate that neural populations at the mesoscale level possess two unrelated timescales, derived from broadband and gamma power signals. They both increase in sleep, especially NREM3, although broadband signals have stronger autocorrelations and longer timescales. Broadband timescales increase ascending the sensorimotor-association hierarchy, while gamma timescales decrease.

### NREM3 slow waves impact timescales at the single-event and global levels

To investigate the physiological phenomena altering timescales in sleep, we turned to NREM3 slow waves, as they constitute a prominent example of an oscillatory process (Nir et al., 2011) (Figure 5). We computed slow waves in a complementary 10-minute dataset of NREM3 sleep with the same channels as the main dataset. For slow waves detection, we used both the low-frequency (0.5-4 Hz), which is closest to slow waves, and the gamma power (30-80 Hz) signals. The latter was necessary to determine the polarity of the slow waves in the bipolar-referenced contacts, following similar procedures as previous detectors (von Ellenrieder et al., 2016). We found a decrease of gamma power around slow waves’ trough, suggesting an accurate slow wave detection (Frauscher et al., 2015) (Figure 5A). Although the slow-wave and gamma signals were time-locked, there was high variability in the gamma power decrease (s.e.m. in Figure 5A), attributable to slow wave event variability and regional differences.

**Figure 5.**
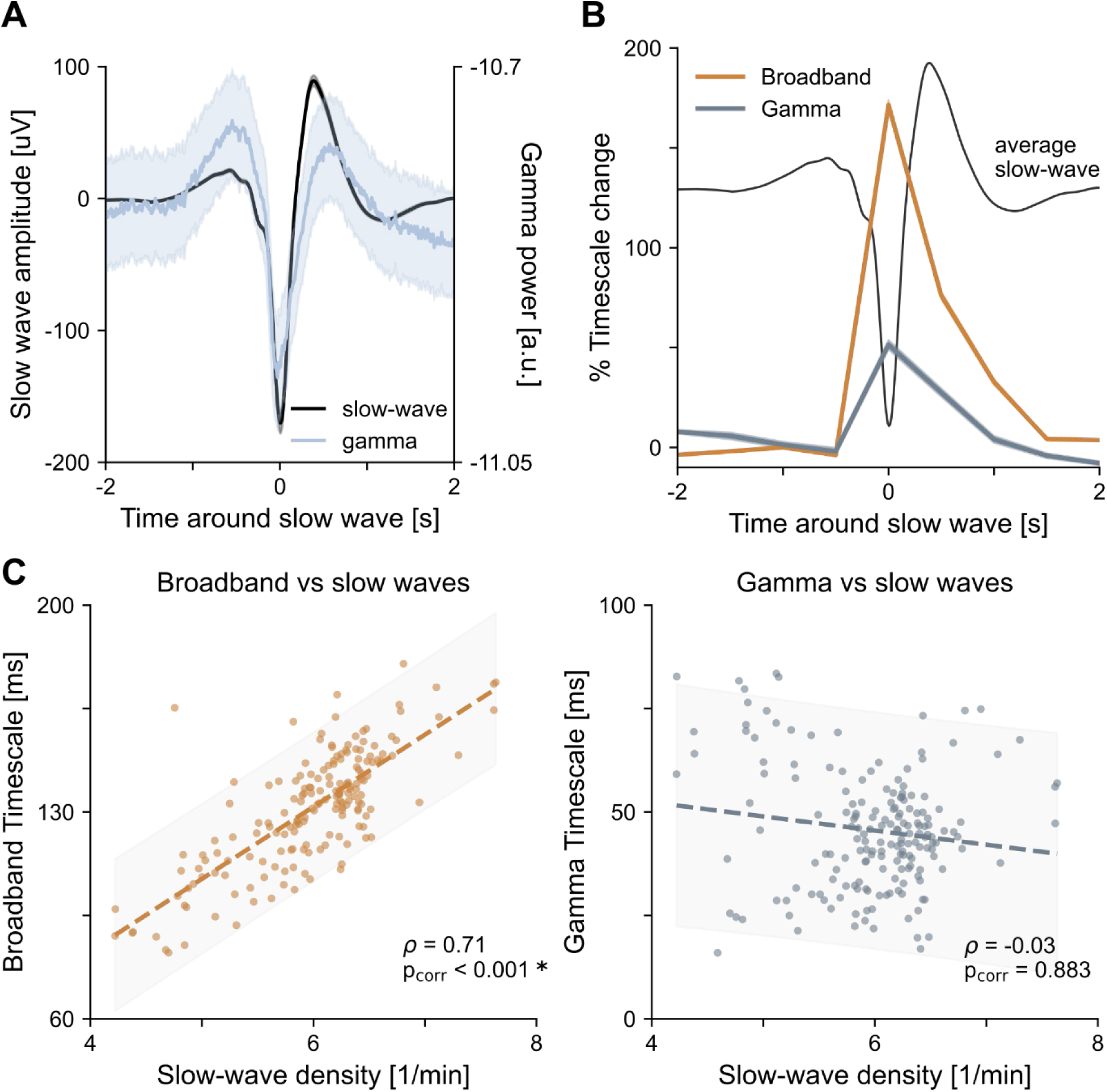
Impact of NREM3 slow waves on broadband and gamma timescales. A) Average profiles of the two signals used to detect intracranial slow waves time-locked to slow-wave troughs. The low-frequency (0.5-4 Hz) “slow-wave” iEEG signal (black) and corresponding gamma power (30-80 Hz, light grey) were used. Averages computed first across channels within a patient and then across patients (n=85121 total slow waves from N=91 patients). Solid lines represent average values, shaded areas the s.e.m. across patients. Note the time-locking between the trough of the slow wave and the decrease in gamma power. For this analysis, a complementary 10-minute dataset with the same NREM3 channels was used. B) Increase in broadband (brown) and gamma (grey) timescales around the trough of slow waves. The average slow-wave profile of panel A is also plotted for visual aid. We computed timescales around slow waves in a sliding-window manner, with 1s windows and 0.5s overlap. Timescale values were normalized to baseline (i.e. the average of −2s and +2s values) such that 0 corresponds to no change. Solid lines represent average values, shaded areas the s.e.m. across the 180 HCP-MMP areas. Note how broadband timescales increase almost two times (171%), while gamma ones only 51%. C) Relationship between timescales and slow-wave density across cortical areas of the HCP-MMP parcellation, for broadband (left) and gamma (right) signals. The correlation is computed with Spearman’s correlation coefficient and p-values are corrected for spatial autocorrelation via permutations with the “vasa” method. Shaded areas represent the 95% prediction interval of the regression. Note the different scales of the y-axes. s.e.m. = standard error of the mean.

After identifying individual slow waves, we linked their occurrence to timescales. Slow-wave occurrence increased both broadband and gamma timescales in a time-resolved manner (Figure 5B). The average percentage change from baseline for broadband timescales reached 171% at the slow-wave trough, while it was only 51% for gamma timescales, or three times lower (Figure 5B). Additionally, at the single-channel level, we found that 99.7% of the channels followed the increasing pattern for broadband timescales, contrary to only 65.2% for gamma ones.

There was also an effect of overall slow-wave density on the timescale value. When correlating slow-wave density, i.e. the number of slow waves per minute, to timescales we found a strong positive effect only for broadband ones (ρ=0.71, p_corr_<0.001), but not for gamma ones (ρ=-0.03, p_corr_=0.88; all computed with Spearman’s coefficient and corrected for spatial autocorrelation with “vasa” method) (Figure 5C). Slow-wave density itself was weakly related to the anatomical hierarchy, such that associative areas tended to have higher density (ρ=-0.33, p_corr_=0.09; correlation with T1w/T2w). To remove this possible confounder, we corrected for this effect by computing a partial correlation with slow-wave density as covariate, and still found the same relationships (Broadband: ρ=0.66, p<0.001; Gamma: ρ=0.07, p=0.36; uncorrected p-values).

This analysis reveals a strong influence of slow waves on the increase of timescales during NREM3 sleep. The effect is prominent for broadband timescales both at the single-event level and at the overall slow-wave density level. For gamma timescales, the effect of timescale increase at the single-event level is more modest and 1/3 of the channels do not follow the increase pattern.

### Spatial correlations are sleep-stage dependent and relate to timescales in a distance-specific way

We next investigated how spatial correlations (SC) manifested across the cortex, and how they related to timescales, which quantify temporal autocorrelations. As a measure of SC, we computed the maximum cross-correlation (CCF) across all pairs of channels in the same patient and hemisphere (wake: 19459; NREM3: 14804; REM: 10364 total pairs; Figure 1D). The SC across distance between pairs of channels displayed an exponential decay profile, for both broadband and gamma signals (Figure 6A, C). An exponential fit of the SC values versus distance revealed that NREM3 possessed the highest long-range SC values, especially in the broadband signal (Figure 6A). As for ACFs, also SC magnitude was lower for gamma signals relative to broadband ones (compare y-axes in Figure 6A and C). We then computed average SC across stages and areas, using the 38 gray-matter areas of the MNI parcellation (Frauscher et al., 2018). Both broadband and gamma SC increased during NREM3 (Broadband: mean difference 0.018, [0.012, 0.022] 95% CI; Gamma: mean difference 0.003, [0.002, 0.004] 95% CI; CIs computed with 9999 bootstraps) and decreased in REM sleep (Broadband: mean difference −0.011, [−0.016, −0.007] 95% CI; Gamma: mean difference −0.001, [−0.003, −0.0005] 95% CI; CIs computed with 9999 bootstraps), relative to wake (Figure 6B, D).

**Figure 6.**
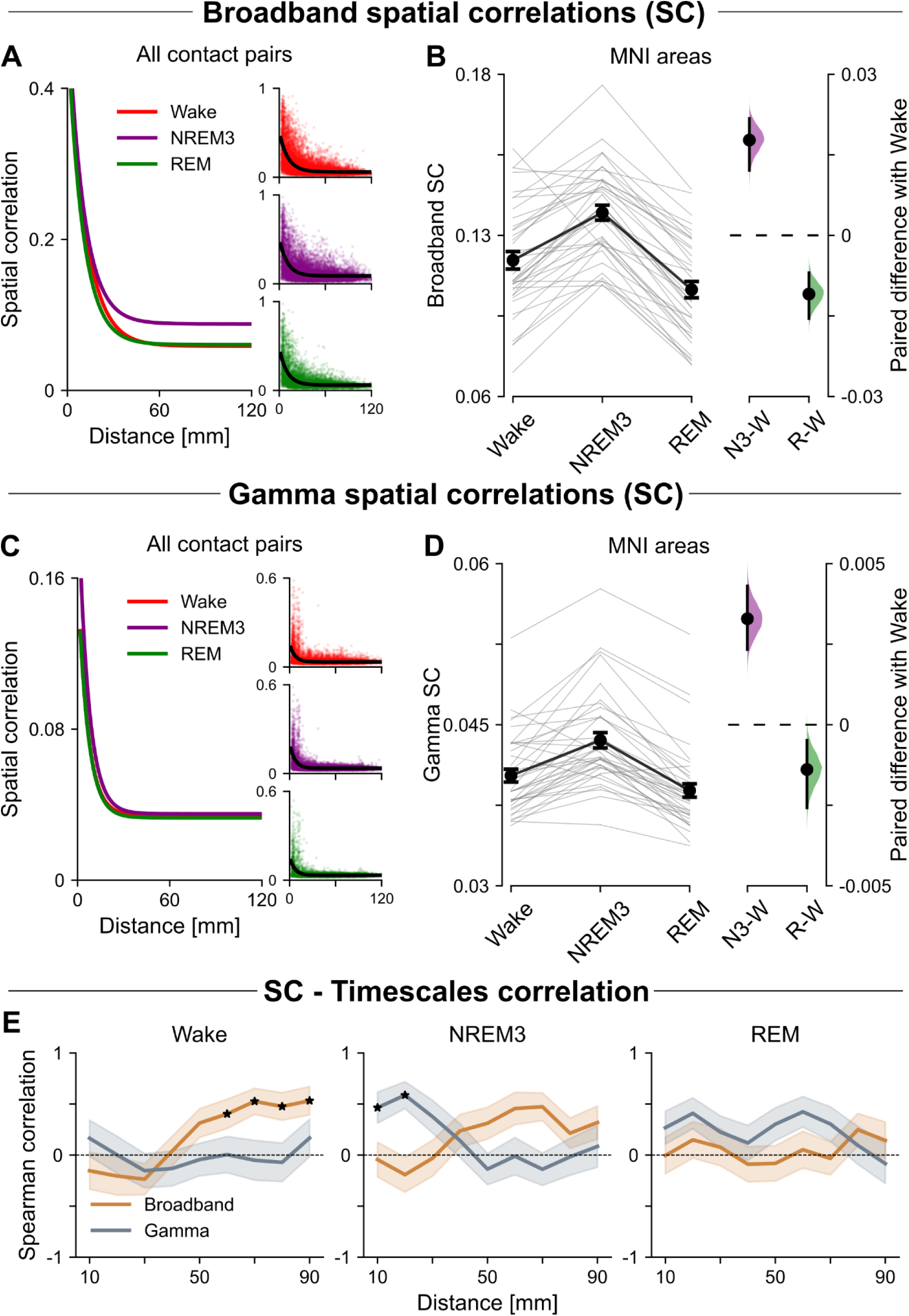
Spatial correlations across stages and their relationship with timescales. A) Broadband spatial correlations (SC) computed from the max CCF between pairs of channels. Across distance, the SC decay like an exponential. The bigger plot shows the fitted curves for wake, NREM3 and REM, while the little insets show the data points and fits for each stage separately; axes labels are the same as for the main plot. For the fits, the SC value for every possible channel pair was used (wake: 19459; NREM3: 14804; REM: 10364 total pairs). Note how SC in NREM3 remain high for long distances. B) Each pair was parcellated into the 38 areas of the MNI parcellation and the average SC was extracted per area with a linear mixed-effects model. Left: Slope plot with the change of broadband SC across MNI areas; thin lines represent each area and black dots average values. Right: Estimation plot with the effect size of the difference between sleep stages and wake. The central dot represents the mean and the vertical bar the 95% confidence interval; distributions are obtained via 9999 bootstraps. C, D) Same as A, B), but for the gamma power signal. Note how correlations are around 5-10 times weaker than for the broadband signal. E) Relationship between SC and timescales, across distances. SC were averaged in 20-mm bins with 10-mm overlap and correlated with timescales for broadband and gamma signals. The correlation is computed with Spearman’s correlation coefficient and p-values are corrected both for spatial autocorrelation via permutations with the “vasa” method and for multiple comparison with FDR correction. Solid lines represent average values, shaded areas the s.e.m. computed from 1000 bootstraps. Stars are distances where the permutation-corrected p-value of the correlation is < 0.05 after FDR correction. s.e.m = standard error of the mean.

Having characterized SC across stages, we finally studied their relationship with timescales across cortical areas. When averaging SC across all distances, they were not correlated with timescales neither in broadband (wake: ρ=-0.07, p_corr_=0.24; NREM3: ρ=0.06, p_corr_=0.50; REM: ρ=-0.003, p_corr_=0.49), nor in gamma signals (wake: ρ=0.07, p_corr_=0.365; NREM3: ρ=0.41, p_corr_=0.006; REM: ρ=0.22, p_corr_=0.09; all computed with Spearman’s coefficient via a linear mixed-effect models accounting for individual patients and corrected for spatial autocorrelation with “vasa” method).

Although SC and timescales seemed to be unrelated at a global level, a more fine-grained analysis considering the distance factor in SC revealed a different picture. For this, we computed the correlation between SC and timescales across distance bins (Figure 6E). When present, the correlation was always positive, indicating that areas with higher timescales also possessed on average higher SC. Wake was characterized by a significant correlation between broadband SC and timescales for long distances (>50 mm; ρ>0.40, p_corr_<0.02). Instead, NREM3 was characterized by a significant correlation between gamma SC and timescales for short distances only (<30 mm; ρ>0.46, p_corr_<0.02) (correlations computed with Spearman’s coefficient, corrected for spatial autocorrelation with “vasa” method and corrected for multiple comparisons with FDR). REM displayed no significant correlation between SC and timescales at any distance (Figure 6E).

These results show that SC are changing in sleep and have a positive correlation with timescales, in a stage and distance-dependent way, with broadband spatiotemporal integration being prominent in wake for long distances and gamma one in NREM3 for short distances.

## Discussion

We found two distinct timescale hierarchies of neural activity recorded intracranially in the human cortex, in the broadband (0.5-80 Hz) and gamma (40-80 Hz) ranges. Both timescales increased in sleep and specifically during slow-wave events. Surprisingly, the two timescales had opposite relationships with the anatomical hierarchy: broadband ones increased along the sensorimotor-association axis in wake and NREM3, while gamma ones decreased and were longest in sensory regions across wake and sleep. Finally, we found that broadband and gamma spatial correlations increased as well during NREM3 sleep and were related to the respective timescales in a distance- and sleep stage-specific way: broadband spatiotemporal integration was present at long distances in wake, while gamma one at short distances in NREM3.

### Broadband and gamma timescales in iEEG

We first report evidence for two distinct neural timescales in the iEEG at the single-channel level. The broadband timescales described here closely relate to previous work on neural timescales in EEG, iEEG and fMRI (Gao et al., 2020; Raut et al., 2020; Zilio et al., 2021). In wake, we reproduced a previous finding on the same dataset, namely that broadband timescales increase along the sensorimotor-association axis (Gao et al., 2020). Our analyses show that this holds true also with a different estimation of timescales, i.e. an exponential fit to the ACF, instead of the power spectrum fit used in (Gao et al., 2020). To this result we add, for the first time, investigations about sleep and gamma timescales. On this last point, we focused on gamma power given its prominent role in inter-area communication, higher cognitive functions, and relationship with neuronal firing (Manning et al., 2009; Nir et al., 2007; Vezoli et al., 2021). Gamma activity has been associated with feedforward processing while lower frequencies, which are included and dominant in broadband activity, with feedback processes (Michalareas et al., 2016). Our results support the presence of multiple timescales in neural populations, as recently found in population spiking activity of macaques’ visual cortex (Zeraati et al., 2023). We add to this finding that two opposing timescale hierarchies can be identified from mesoscopic iEEG signals, and across the human cortex.

Our finding of two timescale hierarchies related to broadband and gamma signals might also explain recent reports challenging the concept of sensory and motor areas as being at the bottom of cortical hierarchies (Cooper et al., 2023; Lurie et al., 2024; A. M. G. Manea et al., 2024), as mostly assumed to date (Gao et al., 2020; Murray et al., 2014). In studies of timescale hierarchies, the experimental modality might be key, as areas directly involved in a task might reorganize their neural activity and hence change timescale (Vinck et al., 2023). Our findings on gamma timescales confirm these observations of an “inverted” hierarchy, possibly due to tight links between BOLD, gamma, and single neuron activity (Fedele et al., 2020). Given the mesoscale nature of iEEG, it’s possible that these two timescales relate to the peculiarity of this recording modality, with its high temporal and spatial resolutions compared to scalp EEG or fMRI.

Our results for opposite broadband – gamma timescale hierarchies call for revising the notion of functional neural hierarchies. The notion of processing hierarchy has so far relied on the idea that fast timescales are more efficient for tracking sensory inputs as they unfold over time (Honey et al., 2012; Pinto et al., 2022), and slow timescales for integrating information over longer intervals (Golesorkhi et al., 2021). Another possibility is that, for an inherently fast signal like gamma, slower timescales in power fluctuations may be more efficient for a reliable tracking of, and response to, external stimuli. Therefore, these two hierarchies may be reflecting the building blocks for different functional purposes and network modes in the human brain.

### Neural timescales in sleep

We describe here, for the first time, how individual cortical areas change their neural timescales from wake to sleep across the human brain, showing that NREM3 has the longest timescales, followed by REM and wake. Across all areas, our results confirm previous reports of a global increase of timescales in sleep, based on scalp EEG and spiking activity in the medial temporal lobe (Hagemann et al., 2022; Zilio et al., 2021). To this literature, we add the spatial resolution of iEEG, showing that this increase is circuit-specific and hierarchy-dependent. Interestingly, we found that both broadband and gamma timescales followed the increasing pattern from wake to REM to NREM sleep, albeit with different spatial distribution and opposite hierarchies for the two measures. Nevertheless, studies on rodent spiking activity (Meisel et al., 2017) and human iEEG gamma timescales (Müller & Meisel, 2023) have shown a decrease in timescales during NREM sleep compared to wake. A first reason for the discrepancy might lie in regional or cross-species differences across studies. A second reason might be methodological. Although we also extracted and quantified gamma timescales, we did so on 1-second epochs, which are shorter than the 2-minute ones used in (Müller & Meisel, 2023). As a result, our gamma timescales range between 10-100 milliseconds rather than seconds, potentially reflecting different facets of the same underlying physiological processes.

Our results are also compatible with the hypothesis that changes in timescales are caused by circuit-specific physiological processes during sleep. We found that single NREM3 slow wave events increase both types of timescales, and especially broadband ones. Moreover, areas with higher slow-wave density were strongly associated with longer broadband timescales. This finding follows naturally as slow waves imply an increase in low frequency activity and therefore in characteristic timing, which is likely the reason why previous studies on scalp EEG report global increases in timescales during sleep (Zilio et al., 2021).

### Neural timescales and spatial correlations

Finally, we computed spatial correlations as a measure of spatial integration and linked them with timescales, which revealed that higher spatial correlations are associated with areas with longer timescales. We interpret the link between these two measures as a proxy for spatiotemporal integration, showing that it is significantly higher at long distances in wake for broadband and at short distances in NREM3 for gamma. Evidence that the temporal structure of brain activity is related to structural and functional connectivity mainly comes from fMRI studies (Baria et al., 2013; Fallon et al., 2020; Shinn et al., 2023) and, recently, iEEG (Müller & Meisel, 2023). During wake, which is a state of high arousal, areas with long broadband timescales show increased spatial correlations. This result might point to the importance of having both high temporal and spatial integration in the same areas, especially for supporting long-range connectivity.

Perhaps surprisingly, we also found evidence of a widespread decrease in spatial correlations during REM and an increase during NREM3, relative to wake. NREM3 is usually described as a state of decreased connectivity (Massimini et al., 2005; Zelmann et al., 2023), while REM as wake-like (Massimini et al., 2010). This finding can be explained by various factors. First, we used the cross-correlation function, which is a functional measure, as opposed to effective ones previously used (Massimini et al., 2005; Zelmann et al., 2023). This measure indicates similarity of time-series signals and is widely used in the iEEG literature (Kramer et al., 2011), but does not prove connectivity per-se and might lead to spurious relationships (Marín García et al., 2013). Additionally, although cross-correlation does not consider specific frequency bands, it is more influenced by low frequencies, which have dominant power. A possibility here could be that the results we observe might be partially explained by cross-frequency coupling between delta and gamma frequencies, which we didn’t explicitly analyze. Second, the intracranial nature of our data might reveal connectivity patterns not usually accessible by scalp EEG, including for instance limbic areas. Third, the relationship between functional connectivity and sleep stage might not be straightforward. For example, a scalp EEG study found that NREM slow waves can provide enhanced windows of communication between brain areas, and this communication can be global and long-range (Niknazar et al., 2022). Finally, although REM has important similarities to wake, it is a physiologically distinct state, associated already with a “disconnection” at the neuronal level between dendrites and soma (Aime et al., 2022). How such a disconnection within individual cells may reflect on brain-wide circuits remains an open question.

### Conclusions

In summary, we studied two distinct types of iEEG neural timescales in the same neural circuits, related to broadband and gamma activity. Both timescales increase in sleep, but show opposite relationships with anatomy, whereby broadband timescales increase along the sensorimotor-association axis, while gamma ones decrease. Single NREM3 slow waves are an important contributor to the increase of timescales in sleep. Finally, areas with increased timescales also show increased spatial correlations, and these are in turn enhanced in NREM3 and diminished in REM. We speculate that broadband and gamma timescales arise from different physiological processes, and that each may serve different functional purposes. Further investigations characterizing the temporal evolution of timescales from wake to sleep in longer recordings are needed to fully elucidate timescales’ manifestation and role in sleep.

## Materials and Methods

### Dataset

We used the openly available MNI Open iEEG dataset (Frauscher et al., 2018; von Ellenrieder et al., 2020), which consists of intracranial recordings on drug-resistant focal epilepsy patients. The dataset contains 1-minute recordings from 106 patients (48 females, 13–62 years old), in wake, rapid-eye movement (REM) sleep, and non-REM (NREM) 2 and 3 sleep. In this study, we focused on wake, NREM3, and REM. For NREM sleep we chose to focus on NREM3 because it exhibits rich slow wave activity. We also computed NREM2 results, and found largely similar patterns to NREM3, albeit with slightly weaker effect sizes. To not overload the manuscript, we therefore focused only on NREM3. The number of overall contacts per stage changes, since sleep was not recorded in all patients (1772 contacts in wake, 1468 in NREM3, 1012 in REM). Additionally, for the slow-wave analysis we used 10-minute recordings during NREM3 (von Ellenrieder et al., 2020).

All preprocessing steps are described in the original data publications (Frauscher et al., 2018; von Ellenrieder et al., 2020). Briefly, all data was resampled to 200 Hz and bandpass filtered between 0.5-80 Hz. Channels and data segments were selected based on strict criteria and were inspected to be devoid of epileptic activity, making non-epileptic tissue suitable for cognitive studies (Johnson et al., 2020). In the original dataset, the 1- and 10-minute recordings were, in some cases, obtained by concatenating different time segments and adding a 2-second buffer of zeros between them. This buffer time was always excluded in our analyses. Finally, the channels were originally localized in 38 grey-matter areas, here called “MNI parcellation”.

### Cortical parcellations

Single-channel estimates of broadband and gamma neural timescales and slow-wave density were parcellated into the 180 parcels (areas) of the left hemisphere of the Human Connectome Project’s Multimodal Parcellation (HCP-MMP) (Glasser et al., 2016). For the projection from MNI space to surface space (“fsaverage”) we used NiBabel (Brett et al., 2024) and MNE-Python (Gramfort et al., 2013) and followed the procedure as in (Gao et al., 2020). Briefly, metrics for each channel were extrapolated to the other voxels with a gaussian weight function 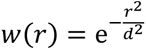 with *d* chosen such that *w*(4 mm)=0.5, i.e. a voxel 4 mm distant weighed 0.5. For every patient, the metric in a voxel was computed as a weighted sum of values from all channels of that patient. Then, the metric values within parcel and patient were averaged, obtaining a parcel-by-patient matrix. Averaging again across patients, weighing by the maximum weight of that patient in the parcel, yielded the final parcel map. The areas with the highest coverage were sensorimotor, temporal, and frontal, but this procedure ensured a smooth map across the whole cortex.

### Anatomical hierarchy (T1w/T2w)

We used the ratio between T1- and T2-weighted structural MRI as a non-invasive, MRI-based measure of anatomical cortical hierarchy. The T1w/T2w is a proxy for myelination and it was found to be high in sensory regions and low in associative ones (Burt et al., 2018). Like in (Gao et al., 2020), we used the group-average myelin map of the WU-Minn HCP S1200 release in the HCP-MMP parcellation, with median values per parcel.

### Neural timescales

Neural timescales were computed per channel on the broadband iEEG and on the gamma power signals. The broadband signal was simply the preprocessed iEEG signal. For the gamma signal, data were first bandpass filtered between 40-80 Hz, then Hilbert transformed, and the power was obtained by squaring the absolute value of the analytic signal and log-transforming it. The log transformation ensured that the data remained approximately normally distributed.

For both signals ACFs were computed in sliding windows of 1 second with 0.5 seconds overlap, using the formula:

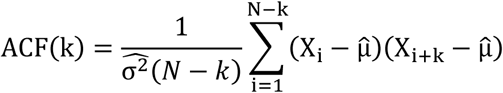

where *k* is the time lag, N the length of the window, and μ̂ and 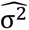1 are the sample mean and variance, respectively. For each channel, ACFs were averaged across windows to yield the final estimate and error. ACFs were computed with the *acf* function from Python’s *statsmodels* (Seabold & Perktold, 2010).

Timescales were then computed by fitting an exponential function f(k) = *a*(*e*^−*k*/τ^ + *b*) and extracting the value of the τ parameter. We already validated this fitting approach in a previous paper (Cusinato et al., 2023) and we saw it provided consistent and robust results, similar to other approaches (see next paragraph and examples in Figure 1C). To validate our results, we compared the broadband timescales in wake with the values reported in (Gao et al., 2020), finding high agreement (ρ=0.74, p_corr_<0.001; computed with Spearman’s coefficient and corrected for spatial autocorrelation with “vasa” method, see later).

For consistency, we validated our results with other approaches that take oscillations into account (Donoghue et al., 2020; Zeraati et al., 2022). Notably, the FOOOF “knee” model yielded similar results in wake, but was more difficult to fit in sleep, due to the presence of long timescales that often resulted in a knee parameter equal to zero, and thus an infinite timescale value. ACFs were fitted between [0, 500] ms for broadband and [15, 300] ms for gamma. We applied a reduced fitting range for gamma because of a fast initial decay that would have otherwise dominated the estimation; and because the ACFs were already quite flat around 300 ms.

### Slow waves detection

Slow waves were automatically detected during NREM3 in the 10-minute recordings, similar to the original publication (von Ellenrieder et al., 2020). Briefly, we used two signals: the low-frequency 0.5-4 Hz slow-wave signal and the 30-80 Hz gamma signal (the filter differs by 10 Hz from the “gamma” used in timescales because this yielded similar but cleaner results).

The detection of slow-wave events was performed separately for every channel, with the following criteria. First, the duration of the positive peak between 0.25-1 seconds, negative trough between 0.25-1 seconds, and a total duration of 0.5-2 seconds in the low-frequency signal. Second, we kept only events with the highest 25% peak-to-peak amplitude, making the selection independent across channels. As a last criterion, we checked whether the average negative trough coincided with a decrease in gamma power (see Figure 5A). If not, we inverted the polarity of the signal and re-computed the slow-wave events, in a similar approach as previous work (von Ellenrieder et al., 2016). Slow waves were computed with the *yasa* package (Vallat & Walker, 2021). Slow-wave density was computed as the number of slow waves per minute. For timescales around slow waves (Figure 5B), we computed ACFs in 1-second windows (0.5 seconds overlap), from −2 to 2 seconds around slow-wave throughs and averaged in the same window across events, finally computing one timescale value per window per channel. We then computed broadband and gamma timescales for each HCP-MMP area at every window following the previous projecting procedure.

### Spatial correlations

As a measure of connectivity between channels, we computed spatial correlations (SC) from the cross-correlation function (CCF) between pairs of channels in the same patient and hemisphere. We used the same hemisphere to avoid increased correlations observed in homologous contralateral regions (Nir et al., 2008). As for ACFs, we computed CCFs for both broadband and gamma signals, in sliding windows of 1 second with 0.5 seconds of overlap, using the formula:

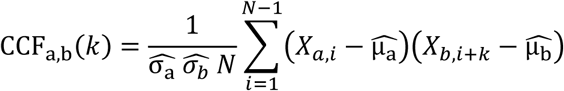

where *a,b are* indices of different channels, *k* the time lag, N the length of the window, and 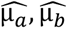 and 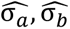 the sample mean and variance of the two channels, respectively (we used *scipy correlate* (Virtanen et al., 2020)). For each pair of channels, the CCFs were averaged across windows to yield the final estimate and error. This yielded a total of 19459 pairs in wake, 14804 in NREM3, and 10364 in REM. Our measure of SC was then defined as the maximum absolute value of the CCF across lags, for every pair of channels (see examples in Figure 1D).

We estimated the decay of SC across distance with an exponential function of the form *f*(*r*) = *a*(*e*^−*r*/*d*^ + *b*), with *r* being the distance between channels (see Figure 6). Finally, we related SC as defined here, and neural timescales. For this comparison, the 38 areas of the MNI parcellation were used, because they provided bigger parcels that allowed for higher pooling of channels. Timescales in this parcellation were estimated with a linear mixed-effects model with patients as random intercepts (*R*’s *nlme* package (Lindstrom & Bates, 1990)). SC values were pooled in a parcel if one of the channels in the pair belonged to the parcel and were also estimated with a linear mixed-effects model. In a second step, timescales were correlated with the mean value of SC in a distance bin of 20 mm, to account for the distance factor in SC; we then slid the bin by 10 mm, going from 10 mm to 120 mm (see Figure 6E). In this way, we correlated timescales and SC across areas at increasing distance, yielding the spatial scale at which the two measures correlated.

### Statistical analyses

For estimating differences between wake and sleep neural timescales across areas we used estimation statistics to show the effect size and distributions, rather than computing a significance test (Cumming & Calin-Jageman, 2024; Ho et al., 2019). Distributions were obtained via 9999 bootstrap samples (*scipy bootstrap* function (Virtanen et al., 2020)) and confidence intervals (CI) from the [2.5, 97.5] percentiles of the bootstrap distribution.

For correlations between maps, we used Spearman’s correlation coefficient, as in previous studies (Gao et al., 2020; Shafiei et al., 2023). Brain maps possess a spatial autocorrelation and vary smoothly across the cortex (Alexander-Bloch et al., 2018; Burt et al., 2020), so the naïve p-values are artificially lowered under the assumption of independent samples. To correct for this factor, a common approach is to generate null maps that preserve the spatial autocorrelation structure but destroy correlations between maps (Gao et al., 2020). We used the “vasa” method (Váša et al., 2018) to generate 1000 permutations of the map on the x-axis and compute surrogate correlations. The p-value was then calculated as the proportion of surrogate correlations bigger than the actual one (two-tailed). For the MNI parcellation, we obtained parcel centroids by pooling parcels from the Destrieux parcellation (Kalamangalam et al., 2021). Then, only cortical parcels were permuted, excluding the hippocampus and amygdala, which were used to compute both the true and permuted correlations. We additionally corrected for multiple comparisons when correlating SC and timescales across distances with the false discovery rate (FDR) method (Benjamini & Hochberg, 1995).

### Data availability

The dataset used is open and accessible at: https://mni-open-ieegatlas.research.mcgill.ca/.

### Code accessibility

We used custom Python code for the analyses, which will be made available on a GitHub repository upon publication.

## Acknowledgments

This work is supported by the Fondation Pierre Mercier pour la science (AT) and the Swiss National Science Foundation (# 320030-227728). The authors thank Dr. Camille Mignardot for helping with brain visualizations.

